# Dynamic Synchronization between Hippocampal Spatial Representations and the Stepping Rhythm

**DOI:** 10.1101/2022.02.23.481357

**Authors:** Abhilasha Joshi, Eric Denovellis, Abhijith Mankili, Yagiz Meneksedag, Thomas Davidson, Anna K. Gillespie, Jennifer Ann Guidera, Demetris Roumis, Loren M. Frank

## Abstract

The hippocampus is a vertebrate brain structure that expresses spatial representations^1^ and is critical for navigation^2,3^. Navigation in turn intricately depends on locomotion; however, current accounts suggest a dissociation between hippocampal spatial representations and the details of locomotor processes. Specifically, the hippocampus is thought to primarily represent higher-order cognitive and locomotor variables like position, speed, and direction of movement^4–7^, while the limb movements that propel the animal are thought to be computed and represented primarily in subcortical circuits, including the spinal cord, brainstem, and cerebellum^8–11^. Whether hippocampal representations are actually decoupled from the detailed structure of locomotor processes remains unknown. To address this question, we simultaneously monitored hippocampal spatial representations and ongoing limb movements underlying locomotion at fast timescales. We found that the forelimb stepping cycle in freely behaving rats is rhythmic and peaks at ~8 Hz during movement, matching the ~8 Hz organization of information processing in the hippocampus during locomotion^12^. We also discovered precisely timed coordination between the time at which the forelimbs touch the ground (‘plant’ times of the stepping cycle) and the hippocampal representation of space. Notably, plant times coincide with hippocampal representations closest to the actual position of the animal, while in-between these plant times, the hippocampal representation progresses towards possible future locations. This synchronization was specifically detectable when animals approached upcoming spatial decisions. Taken together, our results reveal profound and dynamic coordination on a timescale of tens of milliseconds between central cognitive representations and peripheral motor processes. This coordination engages and disengages rapidly in association with cognitive demands and is well suited to support rapid information exchange between cognitive and sensory-motor circuits.

## Main Text

As animals traverse environments, neural population representations in the hippocampus often progress through a sequence of spatial positions, including locations behind, at, and ahead of the animal’s actual position^13–16^. These sequences repeat at ~8 Hz, concurrent with the theta rhythm^12,17^, and are widely thought to reflect a ‘map’^18,19^ of the available navigational space that informs memory-guided behaviors^4,6^. Consistent with this idea, disrupting hippocampal activity or theta impairs subjects’ performance in spatial memory tasks^20–22^, in which correct performance involves locomotion to one or more remembered locations. Thus, hippocampal representations can inform decisions^17,23^ that engage locomotor actions. Conversely, locomotor actions move the animal, and hippocampal spatial representations shift to the new position as animals move.

While interactions between spatial representations and movement are well appreciated, current accounts posit that the processing of information related to spatial representations in the hippocampus is decoupled from the detailed structure of locomotor processes. Specifically, the hippocampus is known to represent higher-order locomotion-related variables, including position, speed, and direction^7^,^24^,^25^, while spinal cord, brainstem, and cerebellum circuits represent and drive specific limb movements^8–11^. The specific coupling of hippocampal representations to limb movements has not been examined, however, and there could be advantages in synchronizing activity across brain systems to facilitate information flow^26^.

We therefore simultaneously monitored neural activity in the dorsal hippocampal CA1 region and the stepping rhythm in rats running on transparent behavior tracks in the context of a spatial memory task. The resulting data included measurements of the frequency of the theta rhythm, the spiking activity of hippocampal neurons including spatially selective ‘place’ cells, and a high-resolution undertrack video from which we extracted rats’ limb movements (**Fig. 1a, Video 1**). Subjects (n = 5) learned and performed a hippocampal-dependent spatial memory task on a W-shaped track^27^,^28^ (**Fig. 1b**). Running trajectories on this task can be classified into outbound (animal running from the center well towards either outer well) or inbound (animal running from either outer well towards the center well), and a correct rewarded sequence corresponds to *Center-Left-Center-Right-Center-Left-Center-Right*, and so on.

**Figure 1:**
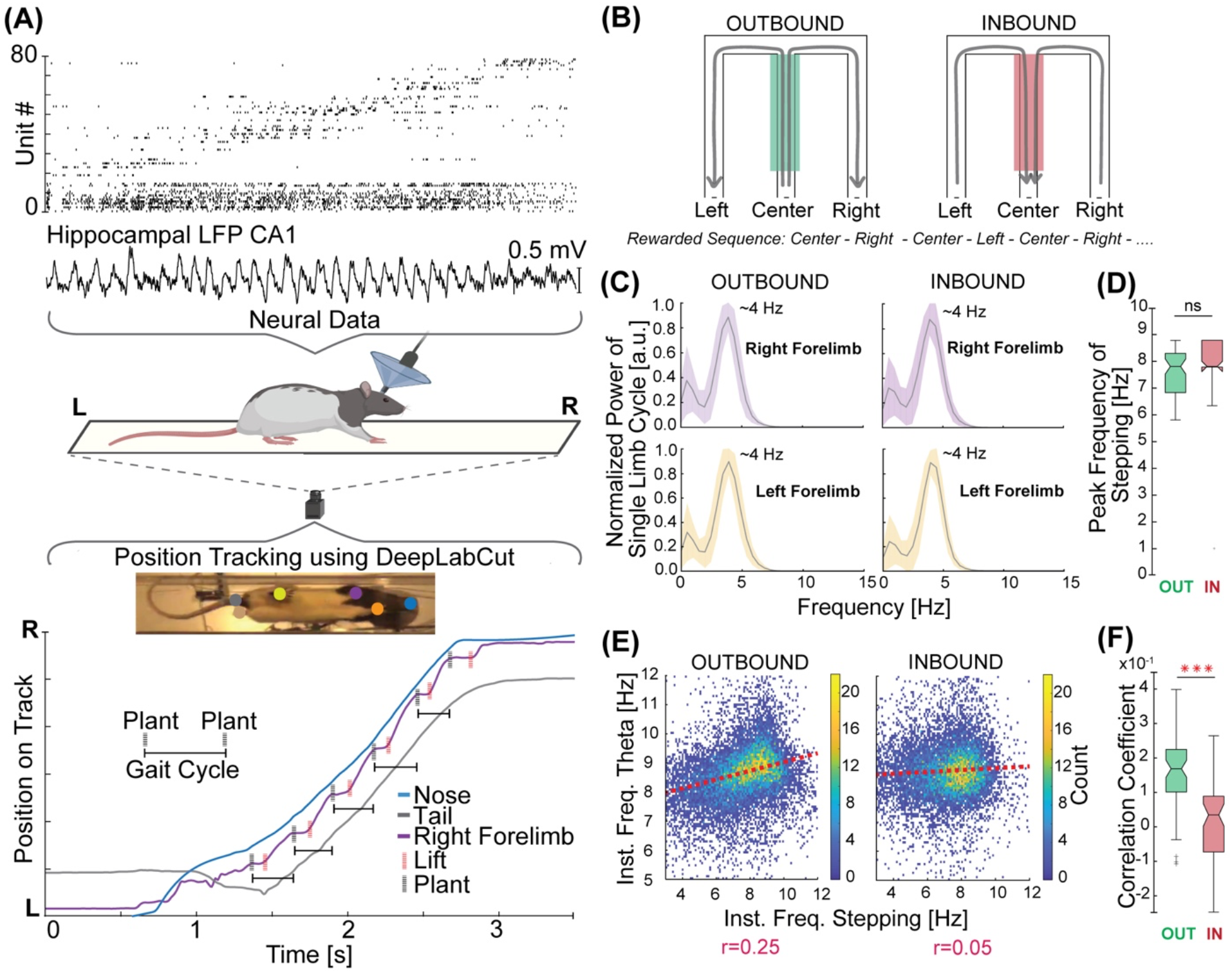
The structure of locomotor activity and its relationship to the hippocampal theta rhythm. **A.** An example spike raster from high-density neural recordings of the rat hippocampus (Rat 1, n=77 neurons) during navigation on a transparent track. For position tracking, a high-speed camera captures the bottom view at 125 frames per second (fps). A machine learning algorithm, DeepLabCut^48^, is trained to track the nose, forelimbs, hindlimbs, and base of the tail of the rat. Simultaneously monitored displacement of the nose, tail and right forelimb. Plant (black dotted vertical lines) and lift (red dotted vertical lines) times of the right forelimb stepping cycle are labelled. **B.** Schematic illustrating the w-track task: Behavioral apparatus and rewarded inbound and outbound trajectories are shown by arrows. The center arm is shaded to signify that the same location is experienced both during inbound and outbound trials. **C.** Power spectral density analysis of the stepping cycle of each forelimb during outbound (*Left*) and inbound (*Right*) traversals of the center arm. Trials for all subjects combined. **D.** Comparison of peak frequency of forelimb stepping observed during traversal of the center portion of the track on outbound (green shaded) and inbound (red shaded) trials (five animals, 61 epochs, outbound, median: 7.8 Hz, interquartile range [IQR]: 6.8 – 8.3 Hz; inbound, median: 7.8 Hz, IQR: 7.8 – 8.9 Hz; outbound vs. inbound Kruskal-Wallis test: p = 0.11, all individual animals p values: N.S. at p = 0.05). **E.** Correlation between instantaneous forelimb stepping frequency and instantaneous hippocampal theta frequency during outbound (*Left*) and inbound (*Right*) runs displayed in binned scatter plots. Color scale corresponds to the count in each bin. Trials for all subjects combined. **F.** The correlation coefficients during outbound trials on the center arm are higher than those during the inbound trials (61 epochs from 5 animals, outbound median: 0.17, IQR: 0.10 – 0.22; inbound median: 0.04, IQR: −0.07 – 0.09, Kruskal-Wallis test, p = 6 x 10^-10^, all individual animals p values < 0.05).

### Dynamic synchronization of hippocampal theta rhythm and locomotor variables

Outbound trials require memory of previous outer arm choices and are more challenging to learn and perform correctly than inbound trials^27^. Additionally, performance on outbound trials is more susceptible to disruption following hippocampal lesions^24^, suggesting that the behavior of the animal has a higher hippocampal dependence for these trials. We therefore compared the relationship between hippocampal neural variables and stepping cycles across outbound and inbound trials. We restricted these analyses to the center arm of the track (30–100 cm, see Methods) as animals approached the T-junction where, on outbound trials, they must choose between the left and right arm.

We first examined the well-known correlation between running speed and the frequency of the theta rhythm^25^,^29^ in that part of the track separately for outbound and inbound trials (n = 61 total recording epochs across five animals). Strikingly, we found this correlation to be stronger on outbound trials as compared to inbound trials (outbound vs. inbound, average difference = 0.14, Kruskal-Wallis test: p = 7.4 x 10^-6^; **Extended Data Fig. 1a, b, f;** comparison to no correlation: outbound, t-test: p = 7.8 x 10^-18^; inbound, t-test: p = 6.2 x 10^-5^). We note that we did not observe a consistent significant correlation between running speed and acceleration on either outbound or inbound trials, in contrast to a recent report^30^ **(Extended Data Fig. 1c, f).**

The differential coupling of movement speed and theta frequency as a function of trial type led us to wonder whether the detailed structure of locomotor processes might also be dynamically coupled to hippocampal theta rhythm, a possibility raised by previous work^31^,^32^. As locomotion consists of cyclic movements of the limbs, we first asked how the overall frequency of these movements compared to the frequency of theta. We found that each forelimb rhythmically moves at a peak frequency of ~4 Hz, together propelling the animal at a stepping frequency of ~8 Hz (**Fig. 1c**). This peak frequency matched the ~8 Hz peak frequency of the theta rhythm and was not different between inbound and outbound trials (5 animals, 61 epochs, average difference = −0.26 Hz, Kruskal-Wallis test: p = 0.11; **Fig. 1d**). Next, we directly assessed whether theta frequency was related to the instantaneous frequency of forelimb stepping and whether this relationship varied by trial type.

Here again we found a trial-type specific coupling. There was a consistent positive correlation between theta and forelimb stepping frequencies on outbound runs (t-test of r values compared to 0: p = 4.8 x 10^-16^; **Fig. 1e, Extended Data Fig. 1f**), but no consistent correlation on inbound runs (t-test of r values compared to 0: p = 0.25); **Extended Data Fig. 1f**). Further, the outbound correlations were significantly larger than the inbound correlations (average difference = 0.14, Wilcoxon signed-rank test: p = 3.3 x 10^-8^; **Fig. 1d**). These relationships could not be explained by differences in running speed (**Extended Data Fig. 1g**). Combined, these results showed that the theta rhythm was more coupled with movement speed and forelimb stepping frequency specifically during the more difficult outbound trials.

### Synchronization of hippocampal spatial representations and stepping on outbound trials

We then asked whether there was also a relationship between stepping and the hippocampal representation of space. Outbound runs on the center arm of the w-track are known to strongly engage theta-paced representations that typically progress, on each cycle, from locations closer to the animal’s actual position toward possible future locations^6^, allowing us to ask whether this progression from current to future is synchronized with stepping.

We used a clusterless decoding algorithm^33^ to determine the location represented by hippocampal spiking activity at high temporal resolution (2 ms time bins, see Methods). Then, we calculated the offset between that estimate of ‘mental position’ and the actual position of the animal to create a distance metric (henceforth, ‘decode-to-animal distance’) that captures the deviation between represented and actual position^34^. We focused on the center region of the track (60–100 cm) on outbound trials, as that region corresponds to the animal approaching the navigational choice point. We then asked whether the decode-to-animal distance was related to the stepping cycle. To measure the relationship with ongoing steps, we used times when the forelimbs touched the track on each cycle (plant times) because these are distinct and identifiable reference points in the stepping cycle and correspond to periods of maximum cutaneous and proprioceptive input from the limbs to the central nervous system^34^,^35^. Here we limited our analyses to those epochs and times where we could reliably decode the hippocampal representation (see Methods).

Strikingly, we found that the plant times of the left and right forelimbs corresponded to hippocampal representations of position close to the actual location of the animal (**Fig. 2a, 2b, 2c; Extended Data Fig. 2a, 2b**). In between these plant times, the hippocampal representation of position typically progressed towards possible future locations and then reset to the actual position of the animal in conjunction with the next forelimb plant (**Video 2**). We focused on theta cycles with appreciable representation of future locations^36^ (see Methods) and quantified this relationship by computing an epoch-wise decode-to-animal distance modulation score (see Methods) that captured the consistency of the synchronization between the hippocampal representations and forelimb plant times. The measured modulation score was greater than the modulation computed from a series of shuffled datasets where the plant times on each trial were shifted by a value chosen from a uniform distribution spanning +/- 70 milliseconds (4 animals, 24 epochs, observed modulation vs. mean of shuffles for each epoch, 60–100 cm on w-track: t-test: p = 1.6 x 10^-8^; **Fig. 2d**).

**Figure 2:**
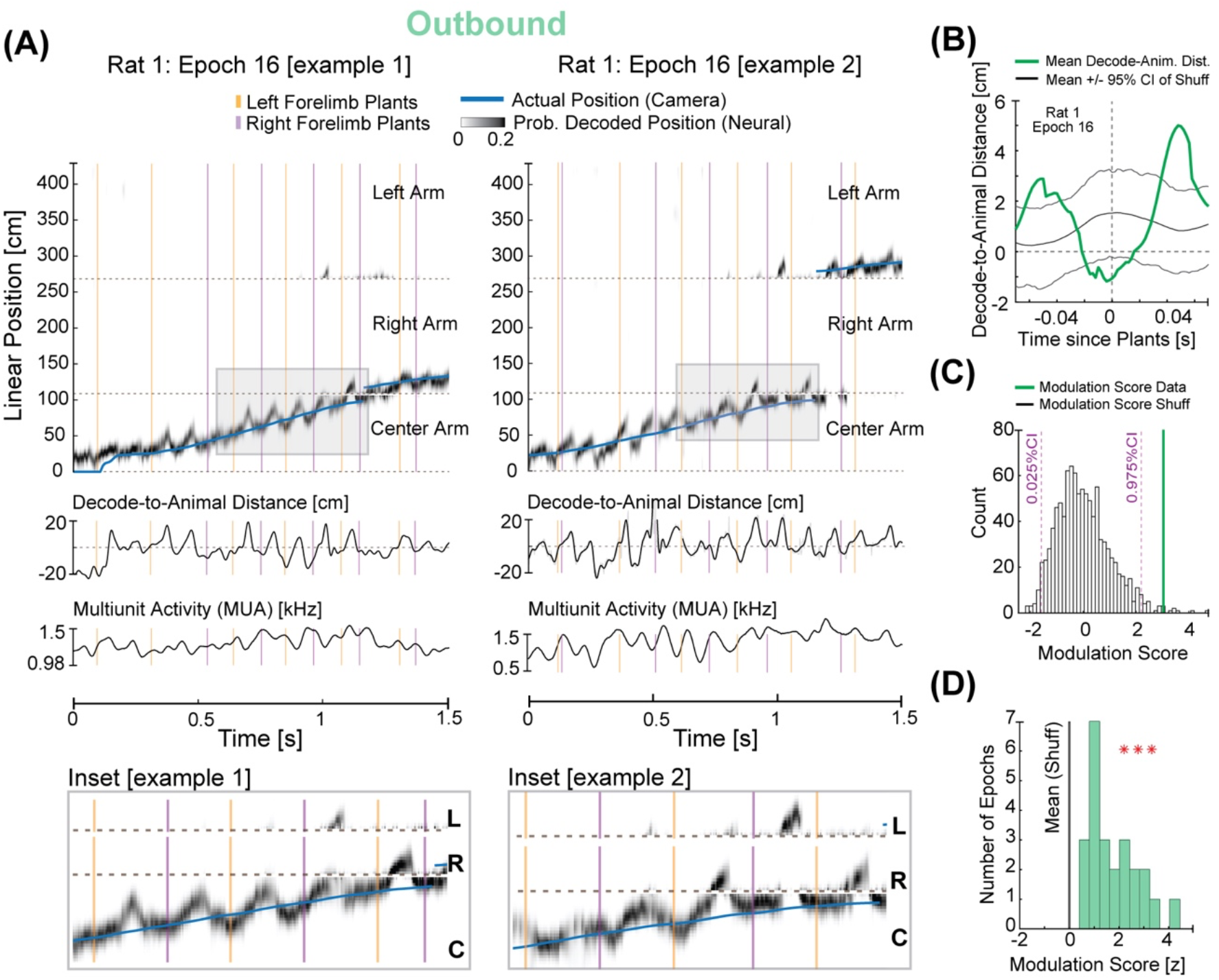
Synchronization between hippocampal spatial representations and forelimb plant times. **A.** Estimation of the represented position based on clusterless decoding: Blue trace represents the linearized position of the rat’s nose. Gray density represents the decoded position of the animal based on spiking. Note that the decoded position can be ahead of, near, or behind the animal’s current position. Orange and purple vertical lines represent the plant times of the left and right forelimb, respectively. Shaded box indicates areas enlarged below. **B.** Decode-to-animal distance trace triggered by forelimb plant times that precede nonlocal representations greater than 10 cm ahead of the animal’s current position for the selected region (60–100 cm) of the center arm (rat 1, epoch 16). Gray lines represent the 95% CI of the shuffled distribution. Dotted line at ‘0’ indicates decode-to-animal distance values corresponding to the current position of the animals’ nose. **C.** Decode-to-animal distance modulation score of the observed data (vertical line, green) and the modulation scores of the shuffled data (bars, gray). **D.** The distribution of decode-to-animal distance modulation score for the observed data in all animals (green, bars) is significantly different from the mean of the modulation score for the shuffled data (black, vertical line; four animals, 24 epochs, t-test: p = 1.6 x 10^-8^, all individual animals p values < 0.05).

This synchronization between plant times and theta sequences also manifested as a synchronization between plant times and the overall multiunit activity (MUA) levels in the hippocampus. In conjunction with rhythmic theta sequences, hippocampal neurons fire rhythmically, such that multiunit firing rates wax and wane on each theta cycle^13^. We computed the degree of modulation of MUA relative to plant times for each epoch and compared it with the mean of the modulation scores for the shuffled distributions (see Methods). As expected from the relationship between plant times and theta sequences, there was a highly significant temporal modulation of MUA and plant times (5 animals, 61 epochs, 60-100 cm on w-track: t-test: p = 3.9 x 10^-7^; **Extended Data Fig. 2c**).

### Synchronization between forelimb plants and hippocampal representation is dynamic

If the coordination between locomotor processes and hippocampal representations is specifically engaged at times of higher cognitive load, we would expect this relationship to be prevalent on outbound trials but not inbound trials. We therefore examined this synchronization during the inbound runs on the center arm of the w-track (**Fig. 3a**, **Extended Data Fig. 3a**, **3b**).

**Figure 3:**
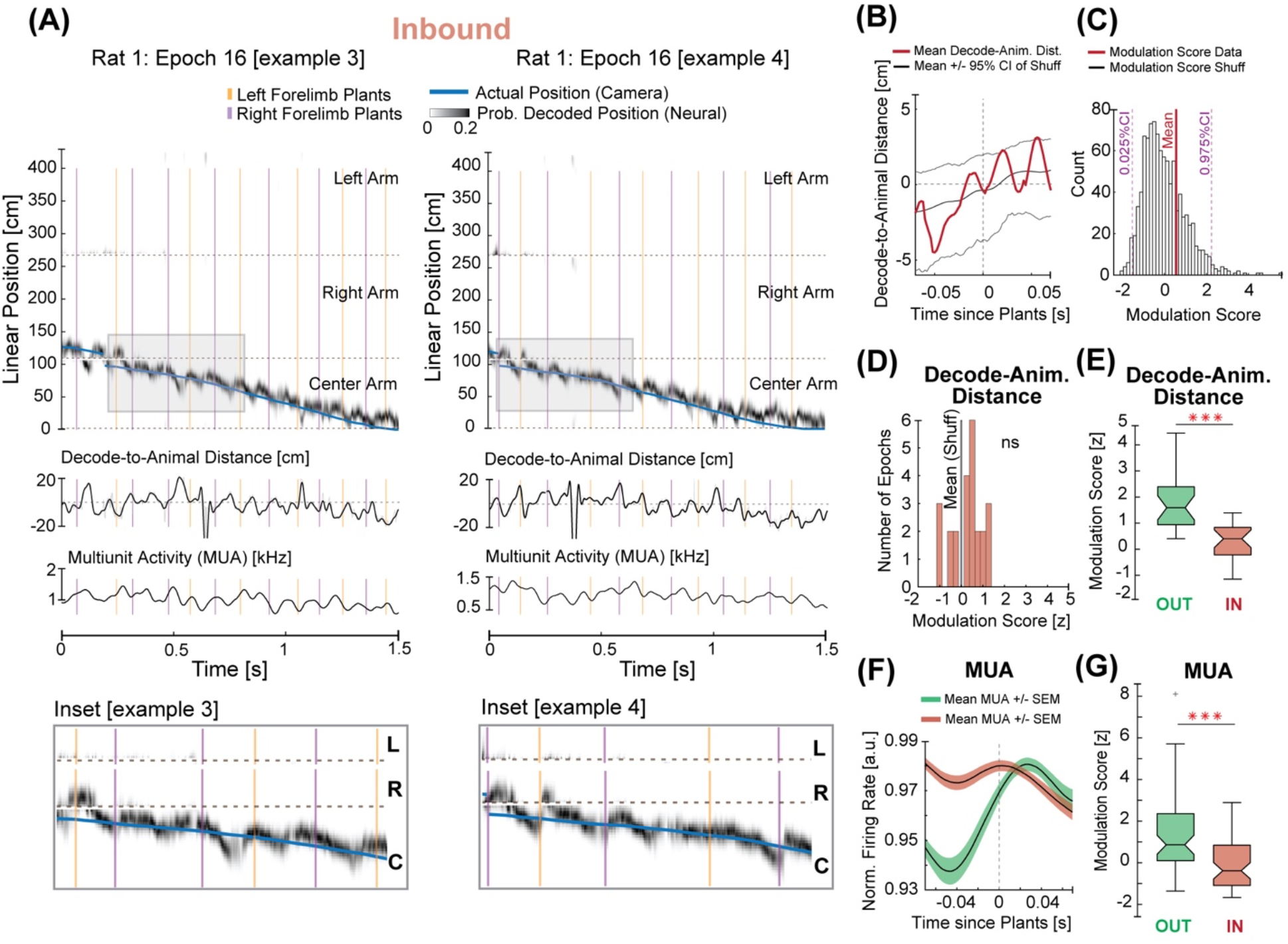
Engagement between hippocampal neural representations and stepping rhythm is task-phase dependent. **A.** Estimation of the representations of position based on clusterless decoding (same as Figure 2) during inbound runs on the center arm of the w-track. Blue trace represents the linearized position of the rat’s nose. Gray density represents the decoded position of the animal based on spiking. Orange and purple vertical lines represent the plant times of the left and right forelimb, respectively. Note that the decode-to-animal distance and multiunit activity rhythmically fluctuate during the inbound runs—shaded regions in zoomed insets below. **B.** Decode-to-animal distance trace triggered by forelimb plant times that precede nonlocal representations greater than 10 cm ahead of the animal’s current position for the selected region of the center arm (60–100 cm) on inbound trials (rat 1, epoch 16). Gray lines represent the 95% CI of the shuffled distribution. Dotted line at 0 indicates decode-to-animal distance values are at the current position of the animals’ nose. **C.** Decode-to-animal distance modulation score of the observed data (red, vertical line) and the histogram of the modulation score for the shuffled distributions (gray, bars). **D.** Distribution of modulation scores for the observed data in all animals (red, bars) and the mean of the modulation score for the shuffled data (gray, vertical line, 4 animals, 24 epochs, t-test: p = 0.08, all individual animals p values: p (rats 1-3): N.S, p (rat 5) = 0.02). **E.** Comparison between the decode-to-animal distance modulation score during outbound (green) and inbound (red) runs on the w-track shows a stronger modulation of decode-to-animal distance by forelimb plants during outbound runs on the center arm (four animals, 24 epochs, Wilcoxon signed-rank test: p = 4.3 x 10^-5^, all individual animals p values: p (rat 1, rat 2, rat 5) <0.05; p (rat 3) = 0.06). ***p < 0.0005. **F.** Modulation of the multiunit activity by the forelimb plant times (mean +/- SEM) during outbound (green) and inbound (red) portions of the w-track. **G.** A comparison of the MUA modulation score during the outbound (green) and inbound (red) runs on the center arm of the w-track shows a more robust modulation during the outbound portions (five animals, 61 epochs, Wilcoxon signed-rank test: p =1.7 x 10^-7^, all individual animals p values: p (rat 1, rat 2, rat 4, rat 5) < 0.05, p (rat 3) = 0.14). ***p < 0.0005.

While we observed clear theta sequences on the inbound runs, we did not observe significant modulation of spatial representations relative to plant times during these periods. This was evident in individual examples (**Fig. 3b**) and in the distribution of the measured decode-to-animal distance modulation scores, which was not consistently different from the respective shuffled distributions (4 animals, 24 epochs, t-test: p = 0.08; **Fig. 3c, 3d**). These inbound scores were significantly smaller than those observed during outbound runs, indicating a low degree of synchronization between the plant times and decode-to-animal distance trace during inbound trials (4 animals, 24 epochs, mean modulation score outbound: 1.75; mean modulation score inbound: 0.27; Wilcoxon signed-rank test: p = 4.3 x 10^-5^; **Fig. 3e**). Similarly, we did not find a significant modulation of hippocampal multiunit activity by forelimb plant times during inbound runs (**Fig. 3f**, normalized mean +/- SEM, **Extended Data Fig. 3c**), and the MUA modulation scores were significantly smaller for inbound as compared to outbound runs (5 animals, 61 epochs, mean modulation score outbound: 1.48; mean modulation score inbound: −0.13, Wilcoxon signed-rank test: p = 1.7 x 10^-7^; **Fig. 3g**). Thus, our data indicate that stepping and hippocampal neural variables are synchronized with each other dynamically according to task-phase in this task.

We then expanded these analyses to other regions of the track, including the outer arms and the regions just past the T-junction (**Extended Data Fig. 4a)**. We reasoned that if the presence of a difficult upcoming choice modulated the synchronization between hippocampal spatial representations and locomotor processes, we would see clear evidence for synchronization on outbound trials before the choice point and little evidence for synchronization past the choice point. Conversely, on inbound trials, we might find evidence for synchronization in the outer arms or T-junction regions, in contrast to the lack of evidence for synchronization in the center arm.

Our results were consistent with those conjectures. We compared the decode-to-animal distance across track regions and found that the most robust modulation was observed during outbound runs on the center arm (Kruskal-Wallis test: p<<0.05, all comparisons on the w-track; **Extended Data Fig. 4b**). We also observed strong modulation of MUA at these times **(Extended Data Fig. 4d)**. Further, we found some evidence for decode-to-animal distance and MUA modulation on inbound runs on the T-junction arm, locations that also preceded a choice (decode-to-animal distance: t-test, p = 0.02; MUA: t-Test, p = 0.04; **Extended Data Fig. 4b-d**).

## Discussion

Our results reveal a remarkable synchronization between ongoing hippocampal spatial representations and the stepping cycle as animals approach upcoming spatial decisions. While previous work showed that various physiological rhythms could be coupled to hippocampal theta rhythm^31^,^32^,^37–41^, we find that ongoing steps are coupled to hippocampal local field potentials, multiunit activity, and the microstructure of spatial representations. This coupling is strongest as animals approach a decision point and synchronized the rhythms such that the hippocampal representation returns to a location close to the animal’s actual position at the time the forelimbs strike the ground.

This dynamic relationship is unlikely to reflect direct drive from sensory inputs to the hippocampus or from the hippocampus to motor outputs. Specifically, the representation of space typically returned to a position close to the actual location of the animal before plant times (**Fig. 2; Extended Data Fig. 2**), and there is no evidence for direct hippocampal output to motor effectors. Instead, we propose that this synchronization reflects a distributed mechanism to coordinate internal hippocampal representations about space (which rhythmically sweep into the future and then return to the animals’ location at theta timescales during behavior) with the ongoing locomotor structure (which provide the strongest sensory signals when the limb strikes the ground^34,35^) such that they concurrently reflect information about the actual position of the animal during plant times. Interestingly, in between consecutive plant times, the hippocampus often represents potential future trajectories. Such an organization^26^ is well suited to segregate information related to environmental sampling^42^ versus planning potential future trajectories^6^ across brain regions involved in fast timescale decision making. Conversely, a lack of synchronization, as on inbound trials in the center arm, may reflect a relative lack of engagement of hippocampal representations in guiding ongoing behavior at these times^27^.

Observations of synchrony between hippocampal local field potentials and sensory-motor processes are not limited to rodents. There is evidence for synchronization between saccades and the hippocampal theta rhythm in non-human primates^37^ and a relationship between button presses and hippocampal theta frequency coherence in humans^43^. Our findings raise the interesting possibility of synchronization between hippocampal representations and movement across species and further suggest that this synchronization would be engaged specifically at times when hippocampal representations are important for storing memories or guiding behavior.

Our data also complement recent work showing that a large proportion of the variance observed in neocortical activity during routine behaviors and in decision-making tasks is related to movement^44,45^. Importantly, however, while those reports identified static relationships on timescales of ~2-5 seconds, we found that locomotor processes are dynamically synchronized with ongoing cognitive representations in the hippocampus on timescales of tens of milliseconds. The existence of these precisely timed representations in the hippocampus, a structure anatomically distant from the sensory-motor periphery, demonstrates widespread coupling between movements, associated sensory inputs, and higher-order cognitive representations.

## Acknowledgments

We would like to thank members of the Frank Lab for helpful discussions, and feedback over the course of the project. The authors are grateful to Daniela A. Astudillo Maya for animal care, Shije Gu for recalculation of per epoch mountain sort metrics, Alison E. Comrie, Brett Mensh, Cristofer Holobetz, David Smith, Eszter Kish, Linda Katona, Michael Coulter, Peter Somogyi, Xulu Sun for commenting on a previous version of the manuscript. This work was supported by the Life Sciences Research Foundation and Howard Hughes Medical Institute grants to A.J. and L.M.F.

## Author Contributions

Design and conception: AJ, LMF; Experiments: AJ, AM; DLC model: AJ, YM; PTP synchronization: TD, AJ; Observation: AJ; Clusterless decoding analysis: ED, AJ; Analysis: AJ, LMF; Shared analysis code: DR, AKG; Surgeries & tetrode adjusting: AKG, JAG, AJ; writing: AJ, LMF; all authors commented on a previous version of the manuscript; Funding: AJ, LMF.

---

**Extended Data Figure 1:**
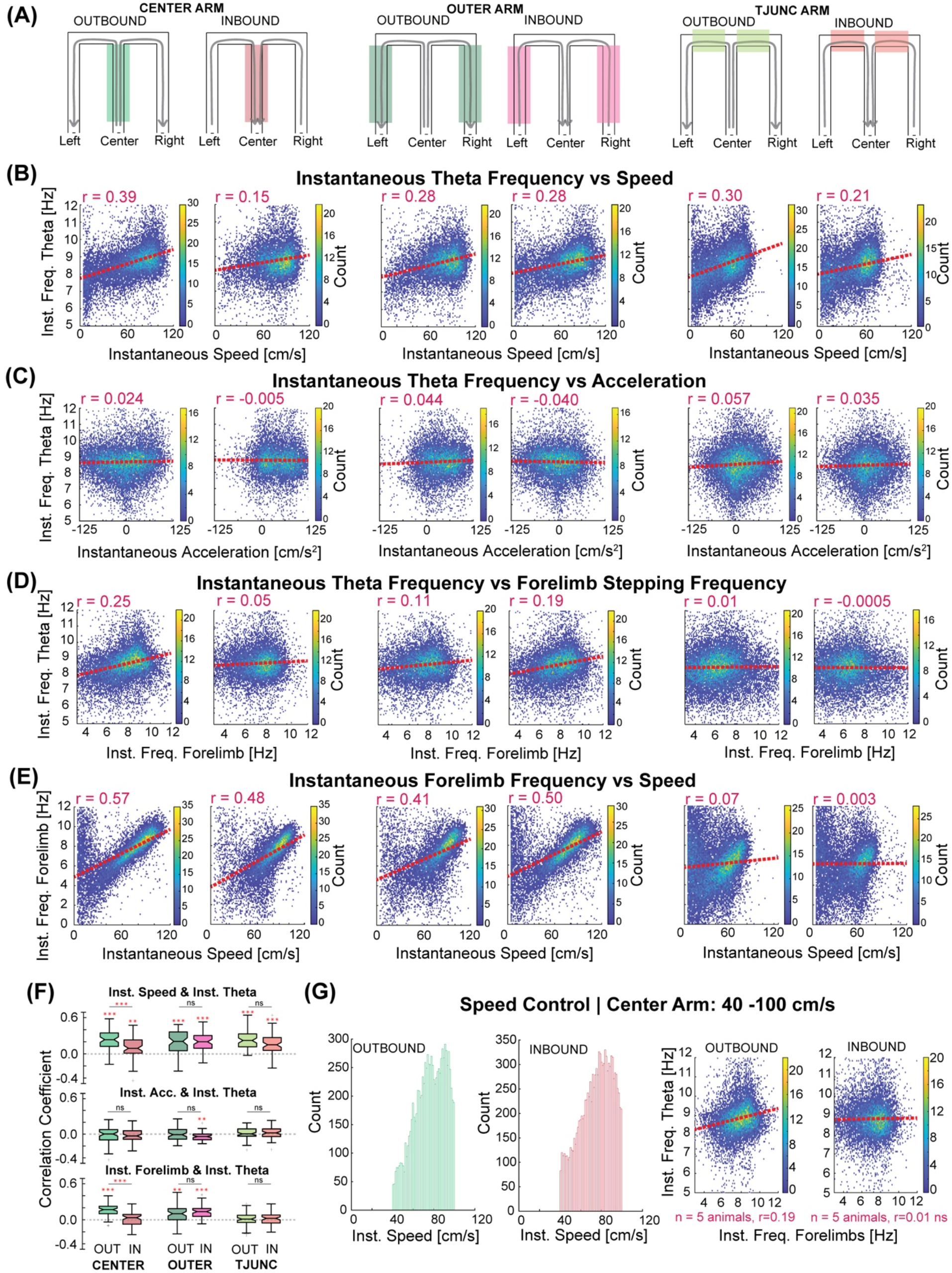
Task-phase specific relationship between hippocampal theta rhythm and the stepping rhythm. **A.** Schematic illustrating the different task phases on the w-track. Gray arrows indicate the direction of movement indicating the trial type (outbound or inbound) in different track regions (shaded boxes). This parcellation defined the 6 task phases on the w-track used in this study: Center Arm – Outbound; Center Arm – Inbound; Outer Arms – Outbound; Outer Arms – Inbound; T-Junction Arms – Outbound; T-Junction Arms – Inbound. **B.** Density plots of the instantaneous hippocampal theta frequency and instantaneous speed on different task-phases on the w-track corresponding to categories in (A). Correlation coefficients (r) for all the data combined are reported on the top left of each panel. Distributions of correlation coefficients computed per epoch for B-E are shown in F. Color corresponds to the count in each bin. Outbound vs. inbound trials on the center arm, average correlation difference = 0.14, Kruskal-Wallis test: p = 7.4 x 10^-6^, individual animal p values: p (rat 1, rat 3, rat 4, rat 5) < 0.05, p (rat 2): not significant (N.S.) at p = 0.05 (comparison to no correlation: outbound, t-test: p = 7.8 x 10^-18^, all individual animals p values < 0.05; inbound, t-test: p = 6.2 x 10^-5^, all individual animals p values: p (rat 2, rat 4, rat 5) < 0.05, p (rat 1, rat 3): N.S.). **C.** Density plots of the instantaneous hippocampal theta frequency and instantaneous acceleration of the animal on different task-phases on the w-track show low correlation coefficients (5 animals, 61 epochs). These variables were not consistently modulated across animals as evidenced by the distribution of correlation coefficients (r, **Extended Data Fig. 1F**) on the center arm during outbound and inbound trials (t-test for outbound values compared to 0, p = 0.41; t-test for inbound values compared to 0, p = 0.42). Color corresponds to the count in each bin. **D.** Density plots of the instantaneous hippocampal theta frequency and instantaneous forelimb stepping frequency on different task-phases on the w-track (5 animals, 61 epochs). Forelimb stepping frequency was strongly correlated with hippocampal theta frequency during outbound trials on the center arm (t-test of r values compared to 0: p = 4.8 x 10^-16^; all individual animals p values < 0.05), and these correlation coefficients were significantly different from those observed during inbound trials on the center arm (Wilcoxon signed-rank test: p = 3.3 x 10^-8^, all individual animals p values < 0.05). Distribution of correlation coefficients for other task phases calculated per epoch is reported in panel F. Color corresponds to the count in each bin. **E.** Density plots of the instantaneous forelimb stepping frequency and instantaneous running speed of the animal on different task-phases on the w-track (5 animals, 61 epochs). Color corresponds to the count in each bin. **F.** Distribution of the correlation coefficients computed per epoch during different task-phases on the w-track. Asterisks (*) indicate that the distribution of correlation coefficients is significantly different from zero (t-test, p < 0.05). Comparisons of the same track-region experienced during outbound and inbound portions on the w-track are highlighted using Wilcoxon signed-rank test. *p < 0.05, **p<0.005, ***p<0.0005. **G.** Running speed control: correlations controlled for the animals’ running speed on the center region of the track during outbound and inbound trials. Analysis was restricted to running speeds of 40-100 cm/s. Histogram of instantaneous speeds during outbound and inbound trials included for analysis (*Left*) and resulting binned scatter plots (*Right*) show that outbound trials on the center arm have higher correlation coefficients compared to those on inbound portions of the center arm.

**Extended Data Figure 2:**
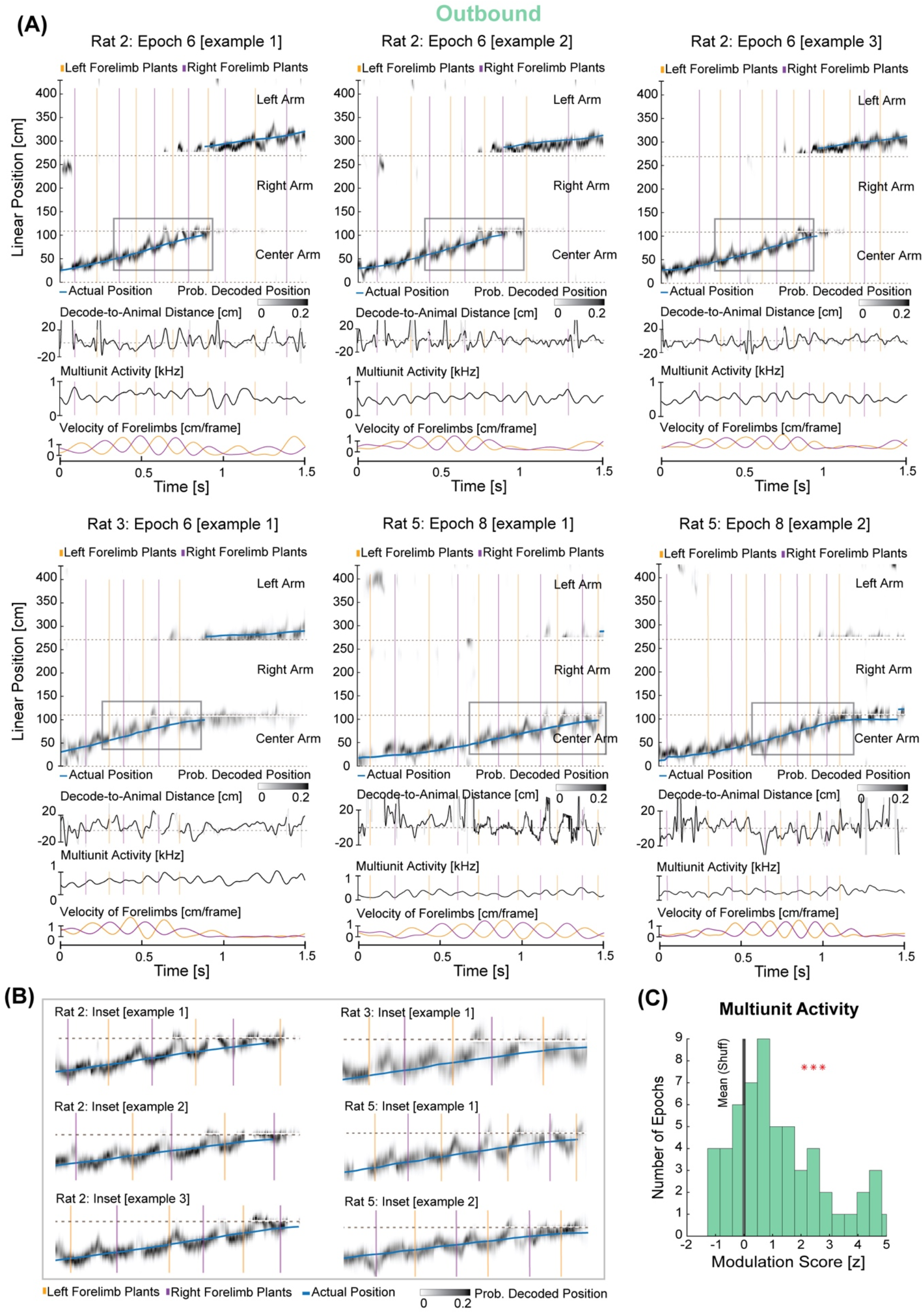
Prominent synchronization of forelimb plant times with neural representation of current position during outbound trials. **A.** Individual examples as in Figure 2 from rat 2, rat 3, and rat 5. Blue trace represents the linearized position of the rat’s nose. Gray density represents the decoded position of the animal based on spiking. Note that the decoded position can be ahead of, near, or behind the animal’s actual position. Orange and purple vertical lines represent the plant times of the left and right forelimb, respectively. **B.** Insets correspond to shaded areas in (A) enlarged to highlight individual examples of the synchronization between hippocampal representations and forelimb plants. Note, forelimb plant times coincide with hippocampal representation of the actual location of the animal. **C.** The distribution of the MUA modulation score for the observed data in all animals (green, bars) is significantly different from the mean of the modulation score for the shuffled data (black, vertical line; five animals, 61 epochs, t-test: p = 3.9 x 10^-07^; all individual animals p values: p (rat 1, rat 4, rat 5) < 0.05, p (rat 2) = 0.07, p (rat 3): N.S.; consistent trends observed in 4/5 animals). Note, rat 3 had 15 electrodes targeted in the hippocampus instead of 30 for rat 1, rat 2, rat 4 & rat 5. ***p<0.0005.

**Extended Data Figure 3:**
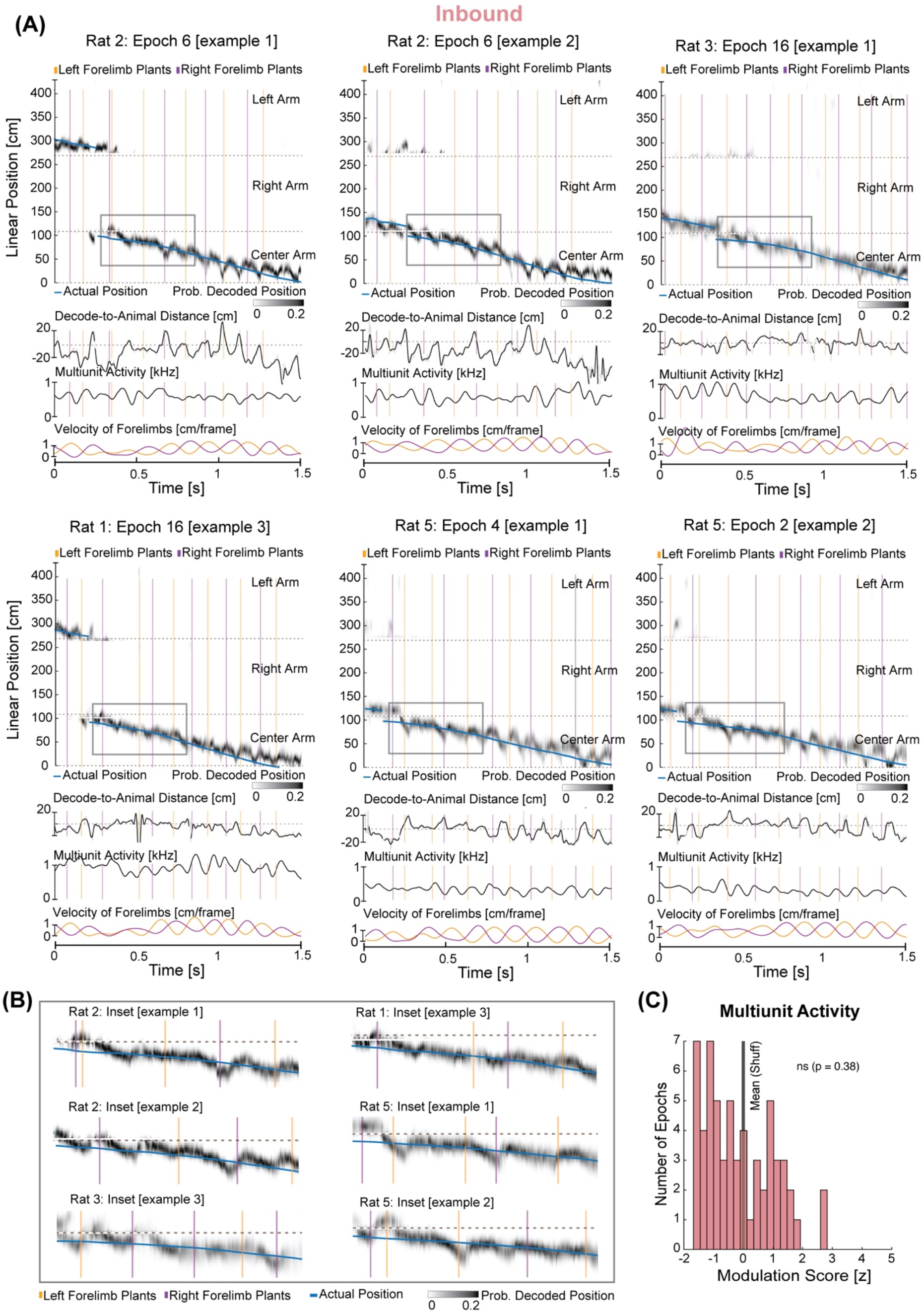
Examples of hippocampal spatial representations and forelimbs during inbound trials. **A.** Individual examples as in Figure 2 from rat 2, rat 3, and rat 5. Blue trace represents the linearized position of the rat’s nose. Gray density represents the decoded position of the animal based on spiking. Orange and purple vertical lines represent the plant times of the left and right forelimb, respectively. Note that the decode-to-animal distance, multiunit activity, and stepping rhythmically fluctuate during the inbound runs—shaded regions in zoomed insets below. **B.** Insets are shaded areas in (A) enlarged to highlight individual examples of hippocampal representations and forelimb plants during inbound trials on the center arm. Note, the lack of coordination between forelimb plants and hippocampal representation during inbound trials. On these trials, forelimb plants could occur when hippocampal decode represents positions that are ahead, concurrent or behind the actual location of the animal. **C.** The distribution of the MUA modulation score for the observed data in all animals (red, bars) is significantly different from the mean of the modulation score for the shuffled data (black, vertical line, five animals, 61 epochs, t-test: p = 0.38, all individual animals p values: p (rats 1–5): N.S.).

**Extended Data Figure 4:**
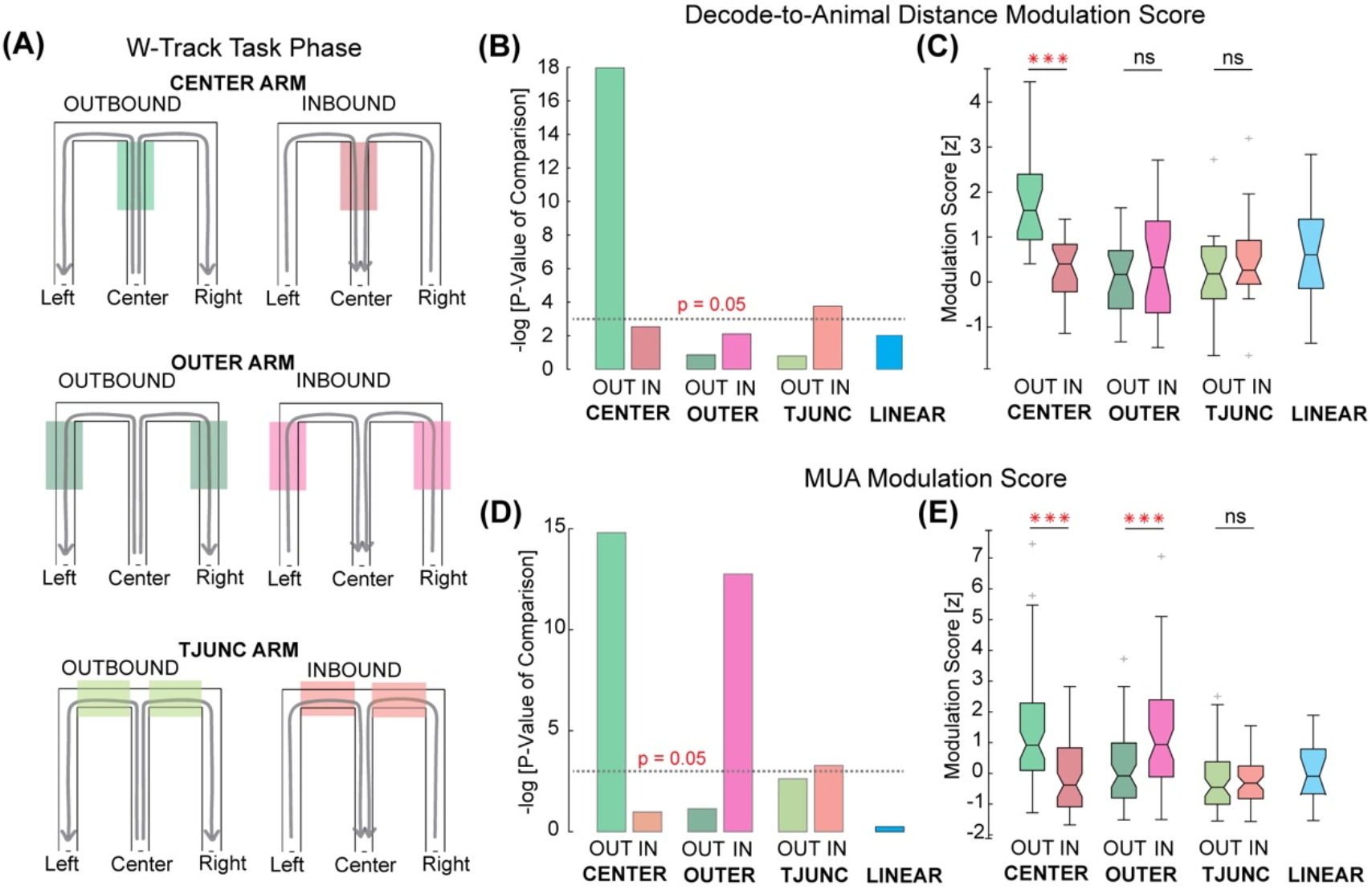
Modulation of decode-to-animal distance and multiunit activity are most prominent during the outbound trials on the center arm of the w-track. **A.** Schematic illustrating the different task phases on the w-track. Shaded portions are highlighted to illustrate regions on the track included for the analysis. **B.** Comparison of the decode-to-animal distance modulation score on different portions of the w-track and a separate linear track. The negative log of the p-value corresponds to the comparison of the modulation score on each portion of the track to that of its shuffled distributions. Dotted line corresponds to p = 0.05 (t-test). Note, a statistically significant decode-to-animal-distance modulation on the T-junction arm during inbound trials (t-test, p = 0.02; all individual animals p values were N.S at p = 0.05). **C.** Boxplots show the distribution of decode-to-animal distance modulation scores calculated per epoch on different portions of the w-track and linear-track. Asterisks (*) indicate that the paired comparisons between outbound and inbound trial types on the same track region were significant (p = 0.05). ***p<0.0005. **D.** Same as **(B)**, but for MUA modulation score. Note, like in the case of decode-to-animal-distance modulation, a statistically significant MUA modulation was observed on inbound trials on the T-junction arms (t-test, p = 0.02; all individual animals p values were N.S at p = 0.05). **E.** Same as **(C)**, but for MUA modulation score. ***p<0.0005.

**Video 1:** Linear track example showing consistent tracking of the nose, tail, and limbs of the rat during movement, first at 1x and then slowed 10 times.

**Video 2:** Forelimb plant times are synchronized with hippocampal neural representations. Green dot is the decoded position of the animal obtained from the firing of hippocampal neurons. The video is slowed down 32 times and pauses briefly at each detected forelimb plant times as an animal is running towards the spatial decision point on the center arm.

## Method Details

### 1. Experimental Model and Subjects

Neural activity (cellular firing and local field potential) was recorded from the CA1 region of the dorsal hippocampus in 5 male Long-Evans rats (*Rattus norvegicus*; 5-9 months old, weighing 500 – 650 g) performing a spatial alternation w-track task^6,27^. Rats were housed in a humidity- and temperature-controlled facility with a 12-hour light-dark cycle. Rats were housed with littermates before experimental manipulation and singly housed in enriched cages during training and food restriction protocols. All experimental procedures were in accordance with the University of California San Francisco Institutional Animal Care and Use Committee and US National Institutes of Health guidelines.

### 2. Behavioral task and neural recordings

Subjects were food deprived to 85% of their baseline weight and pre-trained to run on a linear track for liquid reward (sweetened evaporated milk). This training was done to familiarize the animals with reward wells. After subjects alternated between the two reward wells reliably, they were put back on complete food for at least one week before the implantation surgery. During the surgery, rats were implanted with microdrives^46^ containing 30 (three subjects), 24 (1 subject), or 16 (one subject) independently movable four-wire electrodes targeting the CA1 region of the dorsal hippocampus (all subjects), polymer probes in frontal cortical areas (1 subject) and an optic fiber in the medial septum (1 subject). Only hippocampal data has been analyzed in this study. The hippocampal target electrodes were slowly advanced towards the pyramidal cell layer over 2-3 weeks. Before running on the w-track task (100 cm x 100 cm, track width 10 cm), four subjects also ran on other dynamic foraging tasks in different rooms/contexts. The data presented in this paper is from eight to twenty 15-20 minutes run sessions during learning and performance on the w-track task (number of epochs per animal: rat 1 = 10, rat 2 = 17, rat 3 = 14, rat 4 = 12, rat 5 = 8). The first epoch was excluded from decode-to-animal distance analysis as hippocampal place fields take ~5 minutes to stabilize in a new environment^47^. Each run session was interleaved with 15-20 minutes in an unrewarded rest box. Electrophysiological and video data were acquired using Spike Gadgets hardware and software (https://spikegadgets.com/trodes/). Running trajectories on the w-track were classified into outbound and inbound trials on different track regions, resulting in 6 different task phases during running: Center Outbound, Center Inbound, T-junction Outbound, T-junction Inbound, Outer Outbound, Outer Inbound. Run periods for instantaneous speed and frequency analysis were defined using a velocity threshold (> 4 cm/sec) with a 250 ms buffer. Run periods for decode-to-animal distance and MUA trace modulation analysis were defined using a velocity threshold of greater than ten cm/s with a 250 ms buffer.

### 3. Behavior tracking and monitoring of the stepping cycle

Underfloor video monitoring at 125 frames per second (fps) was performed using wide-angle rectilinear lenses (Theia Technologies; Part Number: SL183M) mounted on AVT Manta cameras (Part Number: AVT-GM-158C-POE-CS; per-frame exposure time: 7.5ms) on both the transparent linear and w-tracks. To ensure that each camera frame was correctly assigned to a corresponding electrophysiological recording time, we captured both the neural data and the position data in a common reference frame using the precision time protocol (PTP). To aid limb identification, the forelimbs of the rats were painted with a white body-paint (https://www.sportsafeproducts.com/) that contrasted with the Long Evans rats’ black hood. Hindlimbs were painted with black body paint to contrast with their white underbellies. A machine learning algorithm, DeepLabCut^48^, was trained to track the distinct body parts of the subjects, including the nose, forelimbs, hindlimbs, and base of the tail. The training dataset included frames from different track portions during various phases of the stepping cycle in both outbound and inbound trials. The model was allowed to run for the maximum number of iterations until its performance reached asymptote. The output comprises of x-y position coordinates for each labelled body part corresponding to each camera frame, along with a likelihood estimate. Position estimates with less than 0.99 likelihood were estimated as the interpolated value of the remaining estimates smoothed with a gaussian window of 0.01 seconds. Velocity of the nose was smoothed with a gaussian filter of 0.15 seconds (filters compensated for group delay). The same model was used to estimate position for all subjects. For position analysis, the nose position was used as the actual position of the rat to correspond closely with previous work that uses an LED on the microdrive for tracking.

### 4. Histology and recording site assignment

In three subjects, the left and right hippocampus was targeted at AP: −4 mm, ML: +/- 2.6 mm, in 1 subject, the left and right hippocampus were targeted at −3.8 AP and +/- 2.6 ML, and in one animal only one hemisphere was targeted at AP: −3.72, ML: +1.26. A screw placed over the cerebellar cortex served as the global reference. Tetrode locations (4 subjects) were marked with electrolytic lesions upon concluding the data acquisition. After a 24hr period to allow for gliosis, subjects were perfused transcardially with 4% paraformaldehyde (PFA). The bottom of the brain was exposed and the brain was left in 4% PFA overnight, following which the tetrodes were moved up, and the rest of the skull was removed. The brain was then transferred to a 30% sucrose solution for 5–7 days, sectioned into 50–100 um slices, and stored in 0.1M PBS-Az. Sections were selected for Nissl staining to enable visualization of the locations of tetrode tips. Electrolytic lesion was not performed for one subject, but all subsequent steps were followed. We used the glial marker Glial fibrillary acidic protein (GFAP) to localize these tetrodes.

### 5. Data analysis

a. **Spike sorting:** Hippocampal spikes were sorted using Mountain Sort (https://github.com/LorenFrankLab/franklab_mountainsort_old)^49^, an automatic clustering algorithm. The output of the algorithm is individual clusters with quality metrics. The quality metrics that were used to plot accepted clusters in Fig. 1a were the Signal to Noise Ratio (SNR>2), Isolation score (>0.90), Noise-overlap (<0.3), and a visual inspection for refractory period violations. Note that sorted spikes were used only for the illustration of spiking activity in Figure 1A.
b. **Power spectral analysis:** Power spectral analysis was performed during run periods using Welch’s method, and each segment was windowed with a hamming window. The result is the power spectral density in each frequency bin (frequency resolution: 1Hz) normalized by the maximum power observed at any bin per epoch. For calculating the peak frequency, we use a minimum peak height of 0.9.
c. **Instantaneous frequency, speed, and acceleration analysis:** Stepping and theta data was filtered (stepping: each forelimb data was smoothed and bandpass filtered between 1 and 6 Hz with roll offs at 0.5 and 8 Hz; theta: hippocampal theta data was bandpass filtered between 6 and 12 Hz with cut offs at 4 and 14 Hz using an acausal filter), Hilbert transformed, and their instantaneous frequency was computed by estimating the average phase difference at each time bin between windows of t-125 and t+125 milliseconds. The instantaneous speed and acceleration were computed similarly in windows t-125 and t+125 milliseconds as the mean of the observed values.
d. **Clusterless Decoding Analysis:**

i. Inputs to the model: We created an encoding model that captured the associations between spike waveform features and the animal’s position at each 2-millisecond time bin as before^33^. The waveform feature used was the peak amplitude of each spike waveform on each of the four channels of the tetrode. Spikes were detected from the 600 Hz – 6 KHz filtered signal when the amplitude on any channel of a tetrode exceeded a 100 μV threshold. The animals’ position was determined by converting the 2D position of the animals’ nose on the W-track to a 1D position based on distance along the track segments (center arm, outer arm, T-junction arm). This linearization is done to speed up the decoding. All trajectories begin with 0 cm representing the center well position, and 15 cm gaps are placed between the center arm, left arm, and right arms in 1D space to prevent the smoothing across adjacent positions from influencing nonoverlapping neighboring segments inappropriately. The code used for linearization can be found at https://github.com/LorenFrankLab/track_linearization.
ii. The model: We used a clusterless state space model (see^33^ for details) to decode the ‘mental position’ of the animal. Decoding used a 20 μV Gaussian smoothing kernel for the spike amplitude features and an 8 cm Gaussian smoothing kernel for position. The state space model had two movement dynamics—continuous and fragmented— which allowed the hippocampal representational trajectory of the animal to move both smoothly and discontinuously through space. This allows us to capture the full range of possible hippocampal spatial representations. The continuous dynamic was modeled by a random walk transition matrix with a 6 cm standard deviation and the fragmented dynamic was modeled by a uniform transition matrix. The probability of staying in either the continuous or fragmented movement dynamic was set to 0.968, which corresponds to 62.5 ms of staying in the same movement dynamic on average, or roughly the duration of half a theta cycle. We have shown the model is insensitive to this choice of parameter^33^. Decoding was done using a causal algorithm with uniform initial conditions for both movement dynamics. A 2 ms time bin and 2.5 cm position bin was used to allow for high resolution decoding. We used 5-fold cross validation for decoding, where we encoded the relationship between waveform features and position on 4/5^ths^ of the data and then decoded the remaining fifth of the data. We repeated this for each fifth of the data.
iii. Outputs of the model:

1. Posterior probability of position: The posterior probability of position is a quantity that indicates the most probable ‘mental’ positions of the animal based on the data. We estimate it by marginalizing the joint probability over the dynamics.
2. Highest posterior density: The highest posterior density (HPD) is a measurement of the spread of the posterior probability at each time bin and is defined as the posterior region that contains the top 50% of the posterior probability values. Using the top values, this measurement of spread is not influenced by multimodal distributions (whereas an alternative measure like the quantiles of the distribution would be). In this manuscript, we use the HPD region size—the total area of the track covered by the 50% HPD region—to evaluate the uncertainty of the posterior probability of position.
3. Decode-to-animal distance: The distance between the decoded position and the actual position of the animal is defined as the shortest path distance between the most likely decoded position (the maximum of the posterior probability of position) and the animal’s position at each 2 ms time bin. The shortest path distance was calculated using Dijikstra’s algorithm^50^ on a graph representation of the track, where the most likely decoded position and the animal’s position were inserted as nodes on this graph.
iv. Epoch inclusion criteria for decode-to-animal distance analysis: For analysis of modulation of the decode-to-animal distance trace around forelimb plant times, we included only those epochs in which we could reliably decode the position across multiple inbound and outbound runs. We estimated this by evaluating a decode quality metric as follows: First, for every run, we computed the mean of the highest posterior density values and the mean of the absolute distance of the decoded position from the current position of the animal. We labelled runs in which either of these values exceeded 50 cm to be ‘noisy’, i.e., where we could not reliably estimate the position of the animal. We then defined the decode noise metric (ranging from 0–1) as the proportion of the length of noisy data to the length of all the data. Those epochs in which the decode noise metric was less than 0.25 for each arm of the w-track, and in which the animal ran each arm at least 10 times were included in the analysis.
e. **Forelimb plant times:** Absolute difference of position data was calculated to obtain instantaneous velocity of each forelimb (i.e., stepping cycle, one value per camera frame). This stepping cycle was then low pass filtered to 6 Hz with a roll-off at 8 Hz to remove outliers and noise events. The stance and swing portions of the stepping cycle correspond to the times when the acceleration of the limb is the minimum and maximum, respectively. An acceleration profile for each limb was created to identify peaks and troughs of stepping rhythm, which was used to define the start and end times of the stance and swing phases. plant times were defined as the midpoint of 10–30% of the stance phase, and lift times were defined as the midpoint of 10–30% of the swing phase. These times correspond to the limbs of the rat fully touching or not touching the track’s surface.
f. **Multiunit activity:** For multiunit activity (MUA) event detection, a histogram of spike counts was constructed using 1.5-millisecond bins; all spikes > 100 μV on tetrodes in the CA1 cell layer were included. The MUA trace was smoothed with a Gaussian kernel (15 millisecond SD).
g. **Decode-to-animal distance/ Multiunit Activity modulation score:** First, we calculated the forelimb plants triggered average of the decode-to-animal distance or multiunit activity trace for each epoch in a time window of ± 70 milliseconds. Then, we computed the modulation score by calculating the sum of absolute deviations from the mean of the observed values in the decode-to-animal distance or MUA triggered trace per epoch. To compare these raw modulation scores across task phases and epochs, we z-scored them using the mean and standard deviation obtained from the null distribution (description below) matched for observed forelimb plants per epoch per animal. All observed plants were included for MUA analysis. For decode-to-animal distance analysis, we included those only sequences that engaged a mental exploration further ahead of the current position of the animal by at least 10 centimeters^36^. Then, each forelimb plant was evaluated in a window of ± 50 milliseconds, and the goodness of the decode-to-animal distance trace was computed in this window by calculating the number of time bins with highest posterior density greater than 50 centimeters. If these values exceeded a total of 10 milliseconds, then those plants were excluded from analysis as we could not reliably estimate the structure of the decoded position adjacent to those plants.
h. **Shuffling analysis:** Plant times were randomly offset between −70 and 70 milliseconds, 5000 times, keeping the inter-event times intact. An event-triggered average of these shuffled times was computed to create the superset of the shuffled distribution data. Then a matched number of events (plants) as observed in the data were randomly selected 1000 times per epoch to create a null distribution of shuffled modulation scores.
i. **Quantification and Statistical Analysis:** All analyses were performed using custom code written in MATLAB 2020a (Mathworks) and Python3.6. Statistical tests used and significance values are provided throughout the text and in figure legends.

